# *Pseudocapillaria tomentosa* Infections in Laboratory Larval and Adult Zebrafish (*Danio rerio*): Development and Advances in an *In Vivo* Anthelmintic Drug Discovery Model

**DOI:** 10.1101/2025.09.22.672915

**Authors:** Connor Leong, Ruby Scanlon, Aisling Kyne, Thomas J Sharpton, Michael L Kent

**Affiliations:** Department of Biomedical Sciences, Oregon State University; Department of Microbiology, Oregon State University; Carlson College of Veterinary Medicine, Oregon State University; Department of Statistics, Oregon State University.

## Abstract

Parasite resistance is an increasing problem in livestock and companion animals. Developing new drug discovery models may improve the identification of novel anthelmintic drugs which will reduce parasite infections. Adult zebrafish (*Danio rerio*) have been previously used as a model for anthelmintic drug discovery by infecting them with the gastrointestinal nematode *Pseudocapillaria tomentosa*. Using larval zebrafish will increase assay sensitivity and throughput because this *in vivo* platform evaluates host parasite interactions and is conducted in multi-well plates. To develop this assay, this study focuses on 1) evaluating infections in 5-30 days post fertilization (dpf) fish, 2) validating the assay with a known anthelmintic used in domestic animals, and 3) documenting the growth and development of *P. tomentosa*. 30 dpf fish had the most robust infections in multi-well plates and the best survival compared to younger larvae. Assay sensitivity was evaluated by aqueously exposing 30 dpf infected zebrafish to emamectin benzoate (a macrocyclic lactone), which successfully reduced infection intensity. *In vitro* larvae hatched from eggs, larval, and adult zebrafish (n=488) were used to document *P. tomentosa* development from 1-37 days post exposure. A change point analysis (CPA) predicted the ecdysis (molting) points as follows (mm): L1/L2 = 0.220, L2/L3 = 0.571, L3/L4 = 1.174, and L4/L5 = 1.584. Identifying the worm molts sizes enables the inclusion of developmental endpoints within trials. Overall, this study will increase the sensitivity of anthelmintic drug discovery because the validated high throughput larval zebrafish model accounts for drug interactions between the host and parasite.

**Graphical Abstract:** 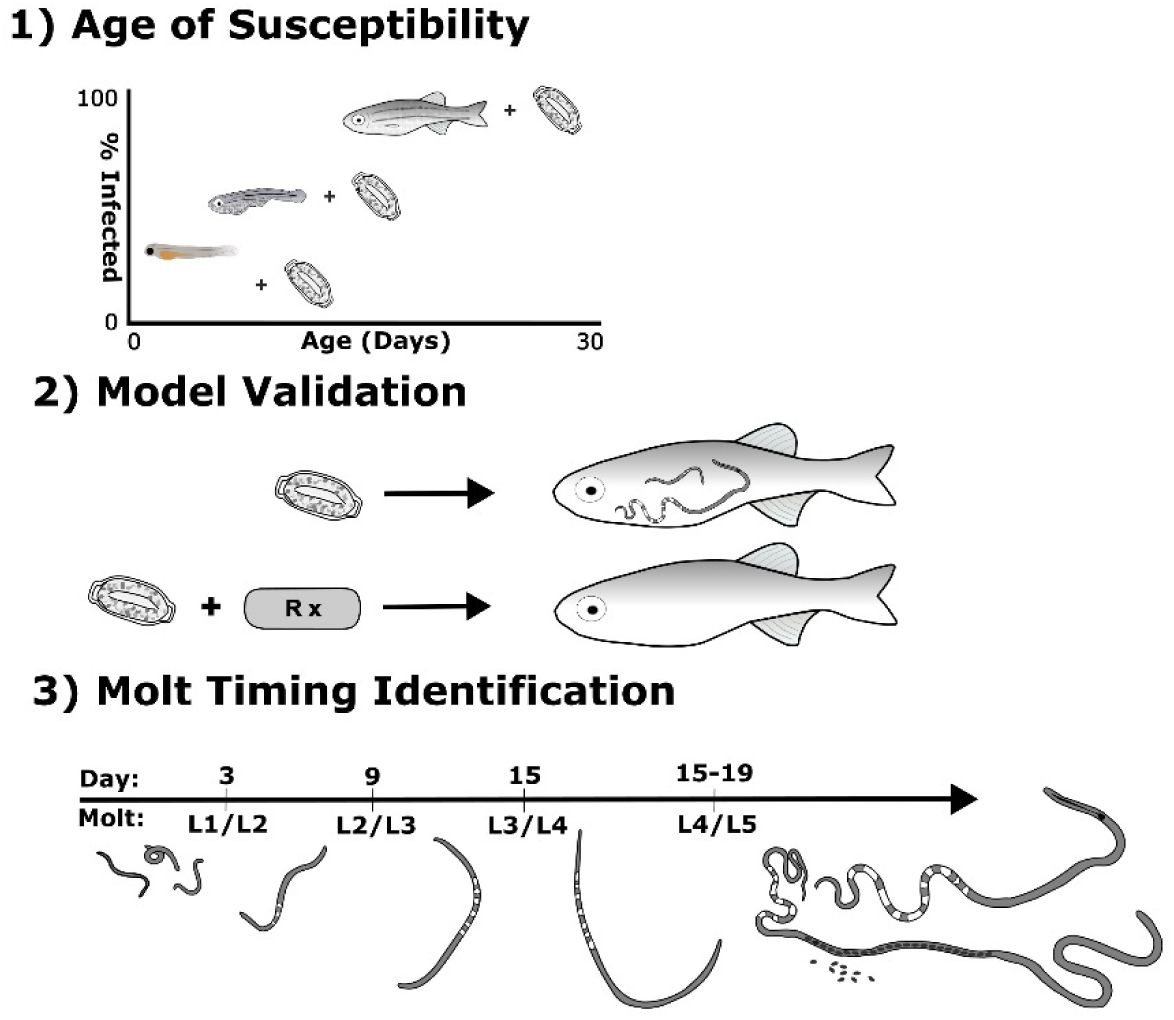

## 1. INTRODUCTION

Nematode parasites present one of the most pressing global health problems for humans and domestic animals, in part due to a limited understanding of etiology and a rapidly growing resistance to anthelmintic drugs. Three of the “13 Neglected Tropical Diseases” listed by WHO are caused by intestinal nematodes as well as three extra-intestinal filarial nematodes. Certain nematodes, such as whipworms of humans (*Trichuris trichura)*, are difficult to treat with available anthelmintics (Adegnika et al., 2015). Moreover, a recently described sister species that also infects humans, *T. incognita*, is highly resistant to two major anthelmintics: ivermectin and albendazole (Venkatesan et al., 2025). Overuse of anthelmintics for nematode parasites has also resulted in a high prevalence of multidrug resistance in trichostrongyles of livestock (Kaplan 2020) and more recently described in hookworm infections in dogs (Jimenez Castro et al., 2019). The rapidly increasing prevalence of drug resistant nematodes demonstrates the pressing need to develop novel anthelmintics for nematodes, which also are important parasites of humans.

No new major classes of ruminant anthelmintics have been introduced in the United States since ivermectin in the 1980’s. The lack of novel therapeutics to combat nematode infections can be attributed to ineffective drug discovery pipelines; modifications to the current pipeline require improvements to effectively prioritize drug leads from large chemical libraries. Early stages of drug discovery are reliant on *in vitro* assays, such as those using *Caenorhabitis elegans* and *Brugia malayi* (Hahnel et al., 2020; Storey et al., 2014). Although capable of quickly assessing large chemical libraries, these *in vitro* approaches do not consider host-parasite interactions that could modulate drug efficacy. In comparison, *in vivo* models enhance drug discovery because they consider the interaction of multiple biological processes such as drug distribution, metabolism, and immune response of the host to the parasite (Carithers, 2017; Vatta et al., 2014). Although mammalian *in vivo* models offer tremendous translational insight, they are low throughput and expensive to implement, limiting the number of drug leads that can be evaluated. Hence, a higher throughput *in vivo* model would be very useful to bridge the gap between *in vitro* and *in vivo* mammalian models. A high throughput *in vivo* model will decrease the likelihood of drug failure in clinical trials due to toxicity, metabolism, and drug distribution because it can quickly screen many drug leads.

Zebrafish are a popular biomedical research model, the second most used animal model behind mice (Kinter et al., 2021). In anthelmintic discovery, zebrafish could likely link *in vitro* and higher vertebrate *in vivo* models (e.g., rodents, sheep) together because of their scalability, cost, and homology compared to mammal models (Patton et al., 2021; Truong et al., 2016). These advantages have been leveraged in other fields to prioritize toxicants that may impact human health (Rericha et al., 2024). First recognized as a pathogen in pond reared carp, cyprinid fishes, and aquarium fishes (Morave et al. 1987; Morave 2001), the intestinal nematode *Pseudocaplliaria tomentosa* is common in many zebrafish research facilities (Kent et al. 2020 b,c). It is associated with morbidity as well a potential cause of non-protocol induced variation when infected fish are used in research (Kent et al., 2012). Zebrafish’s high susceptibility to *P. tomentosa* infections has enabled zebrafish to be used to evaluate both traditional anthelmintics (Collymore et al., 2014; Kent et al., 2019) and novel compounds (Hammer et al., 2024). Hence, the scalability of the zebrafish model and their susceptibility to *P. tomentosa* infections demonstrates their potential application for increasing the sensitivity in early-stage anthelmintic drug discovery.

Infections by *P. tomentosa* show many similarities to gastrointestinal nematodes in mammalian hosts, especially whipworms for which they both are members of the Class Enoplea. Both *P. tomentosa* and whipworms have long-lived infections in which worms penetrate the intestine mucosa and cause severe pathologic changes (Gaulke et al., 2019, 2016; Kent and Sanders, 2020). The *P. tomentosa* life cycle (3-4wk) is initiated by exposing naive zebrafish to larvated parasite eggs and is reliably reproduced in the laboratory (Gaulke et al., 2019, 2016; Kent et al., 2018; Martins et al., 2017). The reproducibility of the infection has enabled us to create a zebrafish infection model to assess the role of the intestinal microbiome during *P. tomentosa* infection (Gaulke et al., 2019), including how water temperature modulates these relationships (Sieler et al., 2025) and the discovery of novel anthelmintic’s associated with and likely produced by gut microbiota (Hammer et al., 2024). While this model system has advanced insight into host-parasite interactions (including the gut microbiome), relying on adult fish for these assays limits the number of drug leads that can be reasonably evaluated.

Hence, the fundamental objective of this study is to improve throughput of the zebrafish infection model by using younger fish, which can be used in preclinical drug discovery. Embryos or larval zebrafish are used with most zebrafish high-throughput model systems, such as those routinely used in toxicology and exposure science (Patton et al., 2021; Truong et al., 2022). Here, we consider whether larval fish could be used to evaluate *P. tomentosa* infection outcomes in a high throughput platform. First, to define this new platform, we investigated the age susceptibility of zebrafish and the patterns of infections by *P. tomentosa* by exposing 5 to 30 days post fertilization (dpf) fish to defined numbers of larvated parasite eggs in multi-well plates. To link the larval fish model to anthelmintic discovery using adult fish, we evaluated the efficacy of a representative macrocyclic lactone, emamectin benzoate. Moreover, while there are at least 20 described species in the genus *Pseudocapilliaria*, no studies provides details on the larval development through the various stages of these worms (Moravec, 2001; Moravec and Justine, 2010; Neyms, 2025). Hence, we documented the development of *P. tomentosa* over 5wk using experimental infections in both larval and adult fish, employing change point analysis to predict molt timing (ecdysis), which can be used as developmental endpoints in the larval fish assay. Validating the larval zebrafish assay and predicting *P. tomentosa* ecdysis provides a sensitive high throughput assay for drug discovery.

## 2. Materials and Methods

### 2.1. Fish Husbandry and Rearing

All research was conducted under the IACUC approval 2022-0280. Wildtype 5D zebrafish were sourced from the Sinnhuber Aquatic Research Laboratory, Oregon State University (OSU), Corvallis, OR, a facility free of *Pseudoloma* and *P. tomentosa*. The Sinnhuber Aquatic Research Laboratory (SARL) follows rigorous and consistent screening protocols using PCR and histology to maintain a specific pathogen free colony (Barton et al., 2016).

### 2.2. Larval Fish

For early stage exposures (fish exposed at 5-8 dpf, Trials I-H-Table 1), zebrafish embryos were obtained by transferring 5 adult zebrafish (3 males and 2 females) into Aquaneering (La Jolla, CA) spawning baskets the evening before spawning. The next morning when lights were turned on the fish were allowed to spawn for 2hr before being returned to their respective tanks. Dead or malformed embryos were removed, and viable embryos were age staged at 6hr post fertilization according to (Kimmel et al., 1995) using a dissecting microscope. At 4 dpf fish were fed Gemma Micro 75 (Skretting, Tooele, UT) two times a day. For late-stage larval fish exposures (fish exposed at 8-30 dpf, Trials I-R), larval fish were obtained from SARL. Once imported to our Nash Hall vivarium at Oregon State University, they were held in static aerated tanks at 28°C and fed 2xs a day with Gemma Micro 75 until transferred to multi-well plates.

**Table 1:**
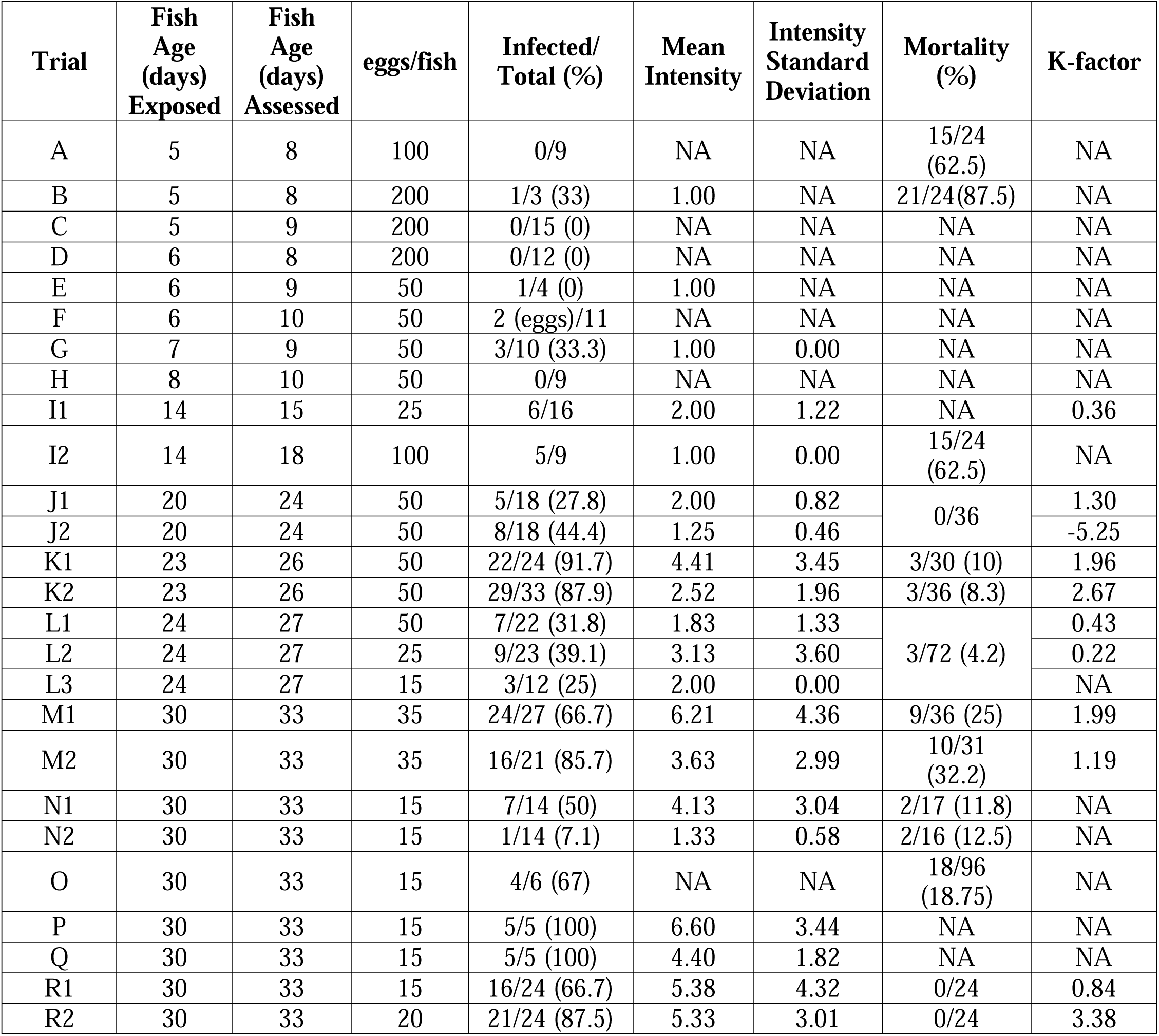
Summary of larval zebrafish trials A-R exposed to *Pseudocapillaria tomentosa*. Trial F, only eggs observed in the intestine of two fish. Trial J2, M2, and N2 represent fish exposed to 0.025, 0.035, or 0.7µM emamectin benzoate respectively. Trial K2 housed triplicate fish in 12 well plates with 1.5 mL of Embryo Medium. Trial L mortality value encompasses all three treatments egg concentrations (L1, L2, L3). Trial M results represented data collected from 3-4 dpe. Trials O-Q are growout trials, and a subset of fish were evaluated at 3 dpe. Trial R represents egg dose response trials. The dispersion (*k*) was calculated for sample sizes greater than 15.

### 2.3. Adult Fish

Adult zebrafish were transferred from SARL to our zebrafish vivarium and were on a flow through system. The Nash Hall Vivarium uses heated city water that is dechlorinated with activated carbon. Temperature within the vivarium was monitored daily (26-28°C) and the following water parameters were monitored weekly, ammonia (0-0.25 ppm), pH (7.6), hardness (0-25 ppm), and conductivity (90-110 uS/cm). Adult fish were fed Gemma Micro 300 5 days per week 1xs a day.

### 2.4. Culturing and Collecting *P. tomentosa* Eggs

A colony of *P. tomentosa* infected fish were used as a source of parasite eggs, which has been maintained in our laboratory for over 15 years (Collymore et al., 2014; Hammer et al., 2024). The infected colony is maintained by cohabiting infected and non-infected fish in a tank on a monthly cycle.

Parasite eggs were collected and purified from infected fish by using a method adapted from (Martins et al., 2017). Approximately 30 *P. tomentosa* infected fish were placed in a 20L spawning tank with wire insert. Fish were housed above the insert for 4 days which prevented the fish from swimming to the bottom of the tank and eating the depurated parasite eggs. Eggs were then concentrated by filtering the collected water through a 200µM screen and collected on a 25µM screen. The filtrate collected on the 25µM screen was transferred into a 15mL conical tube using a transfer pipet (Fisher Scientific: 137117M) and centrifuged at 2000g for 5 min to pellet the eggs. Once pelleted, the supernatant was removed, and 5mL of distilled water was added back into the tube. Purified parasite eggs were larvated in an incubator at 28°C for 5 days. During this time, they were gently agitated with a shaker at speed cycle 2 (Hoefer Red Rotor Shaker, MA). Once larvated, parasite eggs were quantified by first vortexing the eggs and then making 3 wet mount slides. Each wet mount slide contained 20µL of *P. tomentosa* filtrate and was quantified using a Leica DMLB bright field compound microscope at 100x magnification. The mean number of parasite eggs from the three wet mounts was used to calculate the total number of eggs and percent larvated eggs.

### 2.5. Larval Zebrafish Assay Optimization

The optimal larval fish age was determined by exposing 5-30 dpf zebrafish to *P. tomentosa* eggs in several separate trials (Table 1). To optimize fish *P. tomentosa* infections and fish survival various parameters such as egg dose (15-200 eggs/well) and exposure duration (2-4 dpe) were tested for each age stage (Table 1). Fish were housed in 24 well Falcon Plates (product number: 353047) which contained 1mL of embryo medium (EM), comprised of (mM) 7.5 NaCl, 0.25 KCl, 0.5 MgSO_4_, 0.075 Kh_2_P0_4_, CaCl_2_ 0.5, NaHCO_3_ 0.35, and pH 7. The plates were sealed with parafilm, placed on the shaker at speed 2, and held in a 28°C incubator room maintained with a 14:10 light dark cycle. *P. tomentosa* infections were evaluated by euthanizing fish with an overdose of tricaine methane sulfonate (MS-222) (Syndel, Ferndale, WA) then placing the carcass on a glass slide with 50µL of water. A glass coverslip was overlaid, and gentle pressure was applied to flatten the fish to improve visualization of the intestine. Slides were examined at 100x and 200x magnification with a Leica DMLB bright field microscope. More specific assay conditions for each age stage are described below.

### 2.6. 5 dpf fish P. tomentosa Exposure (Trials A-C, Table 1)

At 6h post fertilization embryos were plated in 24 well plates using a disposable plastic pipette (Fisher Scientific, Cat No: 13-711-7M). At 4 dpf the embryos hatched, and the larvae were fed ∼0.5mg of Gemma 75 in the morning. At 5 dpe the EM media was refreshed in the wells and the larvae were exposed to parasite eggs with a *P. tomentosa* egg water concentrate.

#### 2.7. 6 to 8 dpf fish *P. tomentosa* Exposure

(Trials D-H, Table 1), were raised in 24 well plates using the same protocol as the 5 dpf fish. After 5 dpf, the fish were fed ∼0.5mg of gemma 75 every day, with EM media changes every other day. The EM media was changed prior to exposing the fish larvae to parasite eggs with a *P. tomentosa* egg water concentrate.

### 2.8. Greater than 8 dpf fish *P. tomentosa* Exposure (Trials I-R, Table 1)

On the exposure day, a transfer pipet was used to move the fish into 24 well plates. The only deviation to this protocol was trials K1 and K2 which housed fish in 12 well Falcon plates (product number: 353043). Each 12 well plate was filled with 1.5mL of EM/well and the wells in trial K2 contained three fish per well. Across all trials, the water volume in each well was standardized by removing the existing water and then adding the appropriate amount of EM and parasite egg concentrate back to each well.

### 2.9. Larval Fish Grow Out (Table 1, O-Q)

Late stage larval fish (30 dpf) were infected with *P. tomentosa* and grown out to 25 dpe to study nematode development and for predication of molt points. The 30 dpf fish were exposed to *P. tomentosa* in 24 well plates containing 1 fish per well, 1mL of EM media, and 15-30 egg/fish. Plates were sealed with parafilm and incubated at 28°C for 3 days. At 3 dpe, worm burdens were assessed, and then the remaining fish were transferred from their well plates into aerated 2.8L Aquaneering static tanks (48 fish per tank) and maintained at 28°C using water from our vivarium. Fish were fed Gemma 75 one times a day with water changes every 48h. To evaluate worm size and number, fish were periodically euthanized between 3-25 dpe. Wet mounts of the fish carcasses were viewed on a Leica DMLB microscope at X100. The number of adults examined is described in Table 2.

**Table 2:**
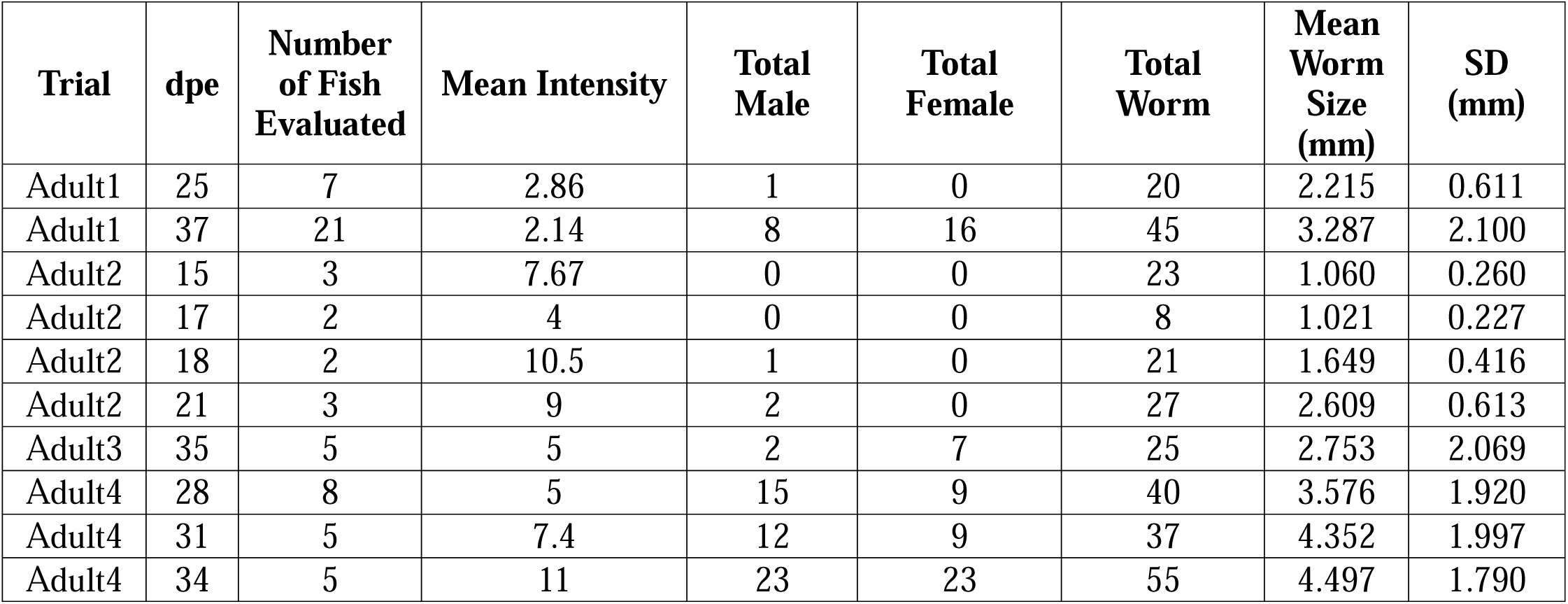
Four Adult fish trials. Fish in Adult1 and Adult4 were exposed to 200 eggs per fish. Fish in Adult2 and Adult3 were infected via a 3 day effluent exposure from a tank with infected zebrafish. Intensity = number of worms/number infected fish.

### 2.10. Adult Fish *P. tomentosa* Exposure

Adult fish were infected with *P. tomentosa* to provide information about late-stage infections because few infected larval fish survived past 10-15 dpe. Fish were held in the Vivarium in 9L Aquaneering tanks and stocked at a density of 30 fish per tank. Fish from trials Adult Trials 1 and 4 (Table 2), were exposed to 200 eggs/fish for 72hr in static water as described in Kent et al (2018). After 72hr, the water flow resumed with standard husbandry practices (Kent et al., 2018, 2019). In comparison, trials Adult 2 and 3 infected fish by exposing them to *P. tomentosa* contaminated effluent from infected fish for 3 days. Following euthanasia, the intestine was dissected from adult fish to better visualize intestinal material in wet mounts.

### 2.11. Nematode Development

*P. tomentosa* development was evaluated between 1-37 dpe, starting with larval worms hatched from eggs, followed by 30 dpf larval and adult fish. Worms were photographed in wet mounts with a Zeiss Axiocam 105 Color camera. The spline function (curve line) on the Zeiss ZEN 3.8 program was used to obtain worm lengths.

### 2.12. *P. tomentosa* Egg Hatching

*In vitro* culturing of *P. tomentosa* eggs enabled the measurement of worms a 24h (L1 stage), the earliest developmental stage. Larvated parasite eggs were placed in 24 well plates containinpg 1mL of hatching medium and incubated at 28°C for 24hr. The hatching medium was composed of 26mL of LB broth, 10mL NCTC -135 (ThermoFisher, Catalog number: 41350026) and 0.942mL of fast grow synthetic bovine serum (MP Biomedical CA, SKU: 0926400-CF). At 24h, the hatched worms were viewed with a Leitz DMBIL inverted microscope.

### 2.13. Histology Processing

Fish at 30 dpf fish were infected with *P. tomentosa* and then collected at 3 dpe for histology to determine if the worm penetrates the intestinal lining as seen in adults. Fish were euthanized with MS-222 and preserved in Dietrich’s. Fish were cut into sagittal sections and the slides stained with hematoxylin and eosin as described by Kent et al. (2020a).

### 2.14. Emamectin Benzoate Toxicity and Efficacy Assessments

Emamectin benzoate (CAS: 155569-91-8) was used to assessed the sensitivity of the larval fish exposure assay because it clears *P. tomentosa* infections in adult zebrafish (Collymore et al., 2014; Kent et al., 2019). Emamectin benzoate powder was obtained from Sigma Aldrich (lot: A0412341). Stock aqueous solutions were made in 100% dimethyl sulfoxide (DMSO, CAS: 67-68-5) and stored in a dark desiccator (humidity < 20%). Serial dilutions in 100% DMSO were made, to obtain the desired concentrations.

The toxicity of emamectin benzoate was evaluated in 10 and 28 dpf fish to identify a safe exposure concentration. All exposures were conducted in 24 well plates filled with 1mL of EM, normalized to 1% DMSO, and had 12 fish per concentration. 10 dpf fish were exposed to (0, 0.0024, 0.05, 1 µM) and 28 dpf fish were exposed to (0, 0.5, 0.7, 1, 10, 100 µM) which were based off previous adult zebrafish emamectin benzoate exposures (Kent et al., 2019). Toxicity was assessed by counting the number of dead and morbid zebrafish with a dissecting microscope at 24hr post exposure.

The toxicity information was used to select the emamectin benzoate concentrations (0.025, 0.035, 0.7µM) for three independent efficacy trials (Trials J, M and N). These trials were conducted with either 23 or 30 dpf fish. Fish were housed in 24 well plates, with 1mL of EM, 1% DMSO, and emamectin benzoate (Table 1, trials: J, M, N). The drug exposed fish and controls were incubated for 3 days before evaluating their infections in the same manner as described in the larval zebrafish assay development section.

### 2.15. Statistics

R studio version 4.3.2 was used for all statistical analysis and data visualization. We applied an array of different statistical analyses, as described below.

The mean intensity and standard deviation were calculated for each trial (adult and larval fish) by first excluding uninfected fish and mortalities. Six worms were not used to evaluate intensity from Trial Adult 1 at 37 dpe (Table 2) because their host at that time point was not recorded. However, their sizes were used in subsequent CPA. The K-factor (*k*) was calculated for larval trials with sample sizes greater than 15 individuals using the following formula 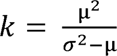.

To conduct the change point analysis, mature females, males, and worms greater than 2.254mm were excluded from the analysis. Moreover, worms from dead fish were also excluded. These worms were excluded from the analysis because sexually mature worms significantly increase in length after they reach the L5 stage. Moreover, it was assumed that immature worms greater than 2.254mm were immature females because they were larger than majority of mature males.

The segmented package was used to perform a bootstrapped change point analysis (CPA) to identify the break points for L1-L5 (Fasola et al., 2018). There were 100 bootstrap iterations where each iteration sampled 80% of the data. The worms in each sampling event were ordered by size before being assessed by the CPA. The CPA used a general linear regression model (worm length ∼ worm order) to predict 4 change points in each random dataset. The size and timing of each molting event was estimated by calculating the mean size and standard deviation of all change point values. If two change point standard deviations overlapped, a t-test (worm length ∼ changepoint values) was used to determine if the standard deviation overlap was significant. These change point values were used to classify the developmental stages of all worms in the dataset. The estimated change point values were cross validated with the appearance of morphological changes (presence of stichocyte bands, spiracles in males and eggs in females), which were noted while measuring the nematodes.

Safe concentrations of emamectin benzoate were selected by calculating the LC50 with the drc r package (Ritz et al., 2015). If the controls had higher than 20% mortality, then the toxicity trial would be rerun. Drug efficacy was assessed by comparing the fish worm burdens between exposure groups using a Wilcoxon rank sum test. Changes in nematode morphology (vacuolation) were evaluated with chi-square analysis of variance. Changes were regarded as significant if the p-value < 0.05. At 9 and 10 dpf, nematode length differences between healthy and moribund fish were evaluated in trial Q1 using an analysis of variance (worm length ∼ dpe*health status).

## 3. RESULTS

Infections were evaluated across 18 trials with larval zebrafish between 5-30 dpf and four trials with adult zebrafish (Table 1,2; Fig. 1-7). Larval worms were clearly observed in the intestinal lumen and possibly the epithelium in wet mounts (Fig. 3). The latter location was confirmed by histopathology, where some worms were clearly observed in the epithelium and associated with epithelial hyperplasia and necrosis (Fig. 2).

**Figure 1:**
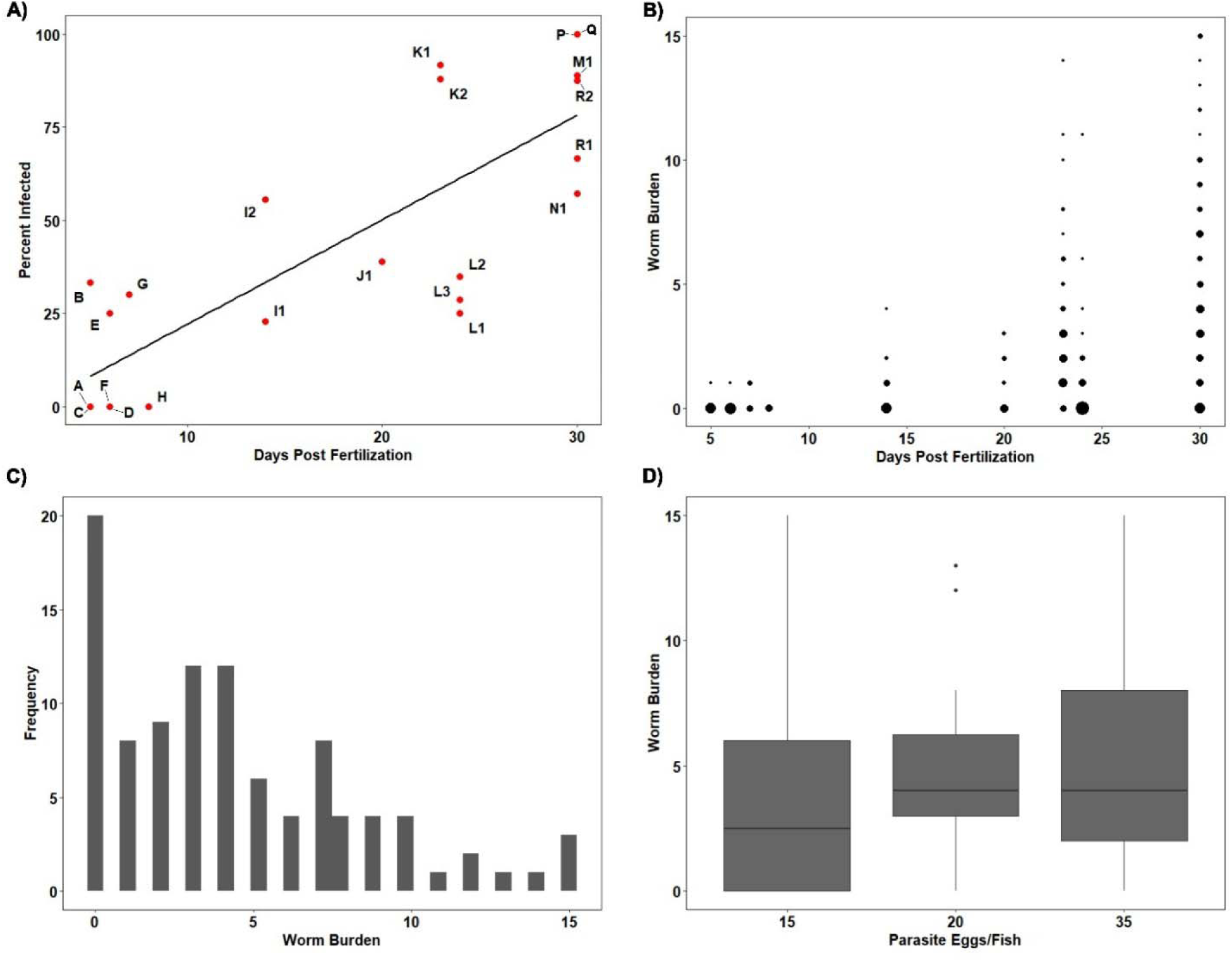
Patterns of *P. tomentosa* infections in larval zebrafish. **A)** Relationship between days post fertilization (fish age) (x axis) vs percent infected (y axis). Red dots represent each individual trial with their letter designation. The black solid line represents a linear regression model lm(days post fertilization ∼ percent infected), showing a significant relationship between days post fertilization and infection success p: <0.001, r^2^: 0.65. **B)** Abundance of worm infections across different fish ages. Y axis is the worm burden of an induvial fish and larger dots indicate a higher number of fish with a specific worm burden. **C)** The worm burden distribution of 30 dpf fish (Trials M, N, O, P, Q, R) which were not exposed to drugs. Fish were exposed to either 15, 20, or 35 eggs/fish and were evaluated by wet mount at 3 dpe. The overall k-value was 1.69 and individual k-values for each trial are reported in Table 1. **D)** 30 dpf fish exposed to 15, 20 or 35 eggs per fish in 24 well plates. At 3 dpe the fish worm abundance was evaluated. Parasite egg dosage had no significant effect on a fish’s worm burden (Kruskal-Wallis worm burden ∼ egg dose, p > 0.05).

**Figure 2.**
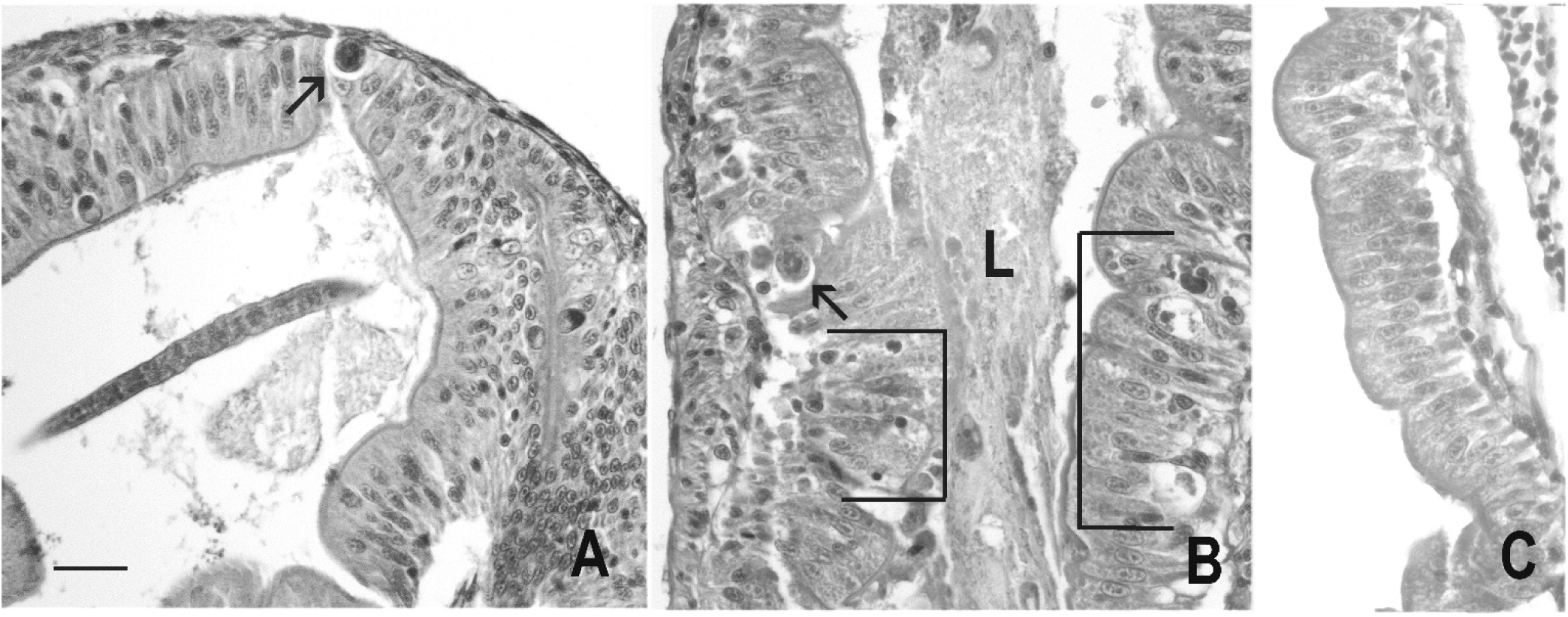
Histological sections of zebrafish larva at 3 days post exposure. Fish was exposed at 30 days post fertilization. H&E. Bar = 20 µm. **A**) Note larva in center of lumen. Cross section of larval worm in epithelium (arrow). **B)** Cross section of worm (arrow). L = intestinal lumen with necrotic debris. Regions of epithelial hyperplasia with intraepithelial necrotic debris are demarked by brackets. **C)** Normal epithelium.

**Figure 3:**
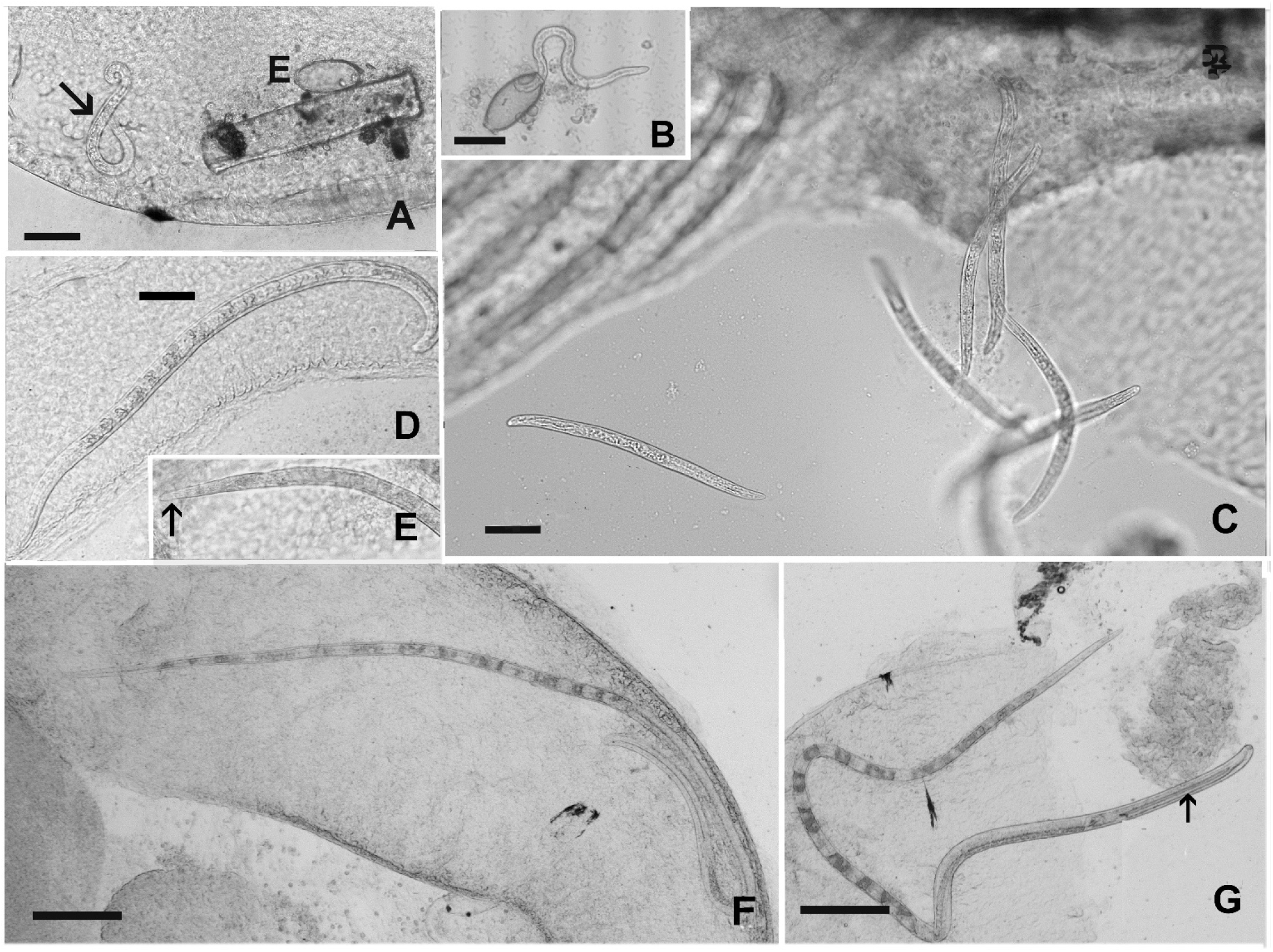
Larval worms in zebrafish intestines a 3-21 days post exposure (dpe) in zebrafish 10 – 30 days post fertilization (dpf). Bar = 50µm unless otherwise indicated. **A)** Fresh hatched larva (arrow) with an empty egg. **B)** Larva emerging from egg after 24hr *in vitro* incubation. **C)** Multiple larvae released from anus of 30 dpf fish examined at 3 dpe. Note, worms are slightly larger and more rigid than larva in A and B. **D)** Worm in at 11 dpe in fish exposed at 30 dpf. Bar = 50µm. **E)** Anterior of L2 stage in 13 dpf fish. **F, G)** Lower magnification of larger worms with bands at from fish at 21 dpe. Bar = 200µm

### 3.1. Fish Age Influences Infection Success

Larval fish (5-8 dpf) (Table 1, Trials A-H) were first evaluated because they are the most common age used in high throughput platforms. The majority of these fish died before evaluation, and the surviving fish had low infection success rates with individuals having no more than one worm (Table 1, Fig. 1A, B). Fish with larger worm burdens were not observed until 14 dpf (Table1, trial I1). Hence, exposing 5-8 dpf fish to *P. tomentosa* did not produce robust infections with adequate fish survival.

In contrast, trials with older fish between 20 to 30 dpf, consistently yielded high prevalence of infections (Table 1 trial J-R, Fig 1). Combining the six 30 dpf trials (M1, N1, P, Q, R1,2) showed a range of 57-100% prevalence of infection, a mean intensity of 5.506 worms/fish amongst the trials, and some fish showing as many as 15 worms/fish (Fig. 2a). In addition, 30 dpf larval fish had acceptable survival over the 3 day exposure in the 24 well plates, with a mean mortality of 20.05% across all 30 dpf trials (Table 1). Overdispersion (aggregation), reported as *k* value, was observed across the trials, and was more pronounced in older fish (Table 1). Combining data across trials for the 30 dpf fish showed a prominent overdispersion, with a *k* value of 1.694 (Fig. 2C).

### 3.2. Optimized Larval Fish Exposure Paradigm

After testing various assay conditions and fish ages the optimized larval fish exposure conditions are summarized and defined as follows. Larval fish at 29 dpf were obtained from SARL and acclimated to our vivarium in Nash Hall for 24hr. The next day, fish were fed Gemma 75 1hr before being plated into 24 well plates (1 fish per well). Plates are filled with 1mL of EM media, 15-30 eggs per well, and sealed with parafilm. Sealed plates are placed on a shaker (low to moderate shaking) and held at 28°C for 72hr. After 72hr the fish are euthanized with MS-222 before assessing their infection on wet mount slides.

### 3.3. Estimating Nematode Ecdysis (L1-L5)

We followed length and other developmental changes of *P. tomentosa* using three platforms; an *in vitro* worm hatching assay, larval and adult fish assays, providing data for worms development from hatching to 37 dpe. The *in vitro* assay allowed for measurement of one day old worms (n= 15 worms). Larval fish exposures evaluated worm length and morphology between 3-25 dpe (grow-out trials O-Q, n = 172), providing data on worm development from early to intermediate age. The adult fish platform was used to obtain data from intermediate to late-stage developmental periods between 15-37 dpe. A total of 301 worms were evaluated in adult fish, which included mature worms, from four separate adult fish trials (Table 2).

As time progressed following exposure, worms progressively increased in length, exhibited banding of stichocytes starting at 9 dpe, and ultimately maturation of male and female worms. Photomicrographs these worm phenotypes are displayed in Fig. 3 and 4. A bootstrapped change point analysis (CPA), with removal of males (identified by presence of spicules), gravid females, and immature worms larger than 2.254mm predicted the timing and worm length for each larval stage (Fig. 5,6). The mean and standard deviation change point values predictions for each iteration were as follows (mm): L1/L2 = 0.220±0.05, L2/L3 = 0.571±0.127, L3/L4 = 1.174±0.266, and L4/L5 = 1.584±0.225. Although there was an overlap between the L3/L4 and L4/L5 change points there was still a significant difference between standard deviations (p = 1.77 X 10^-24^, t-test).

**Figure 4:**
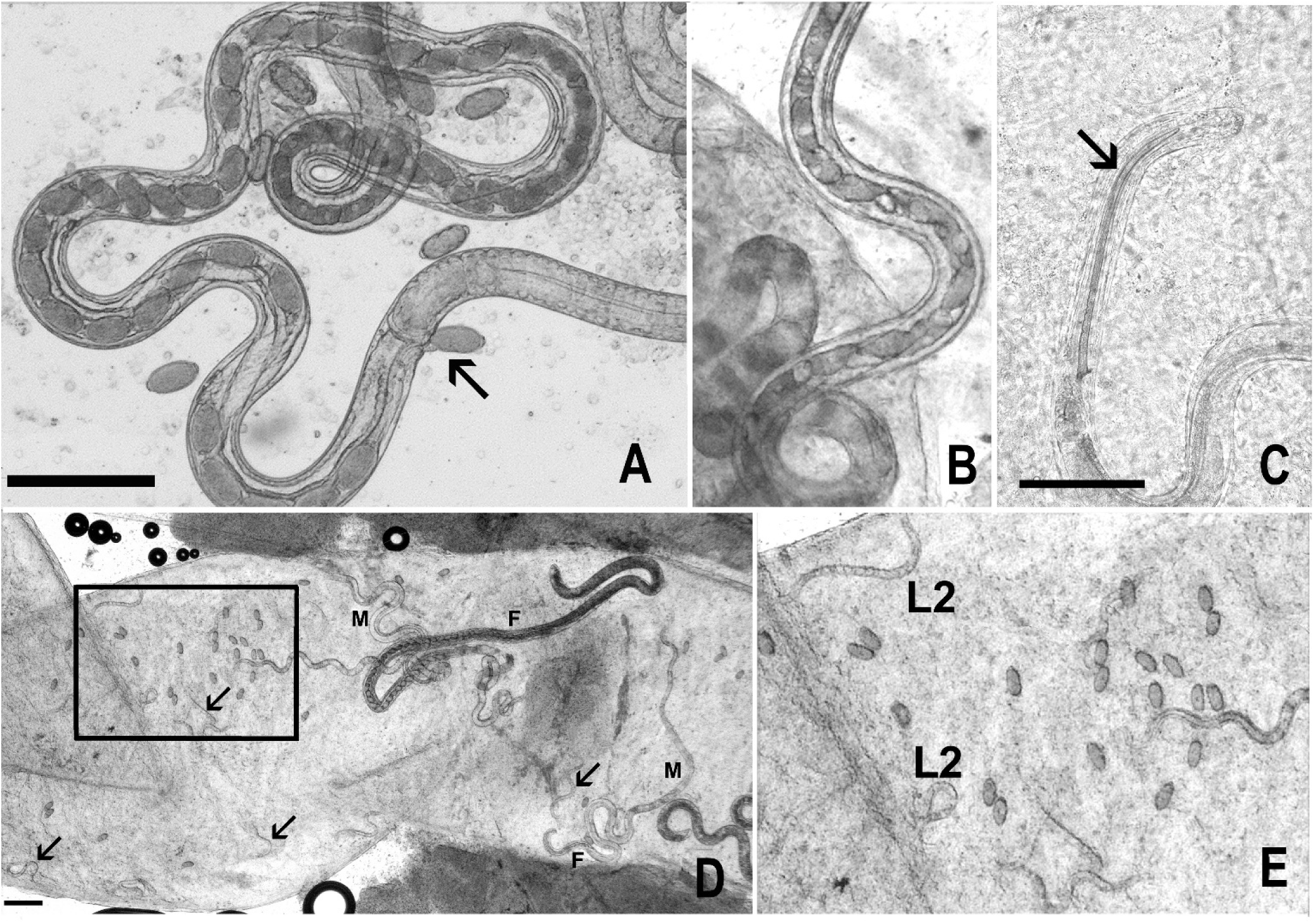
*P. tomentosa* in zebrafish **A**) Gravid female from adult fish at 35 dpe (Trial Adult 3). Bar = 200µm. **B)** Female worm with developing eggs at 25 dpe from larval fish grow out trial (Trial Q). **C)** Male worm in adult zebrafish at 35 dpe. Arrow indicates a spicule in male worm. **D)** *P. tomentosa* L2 stages (reinfection) and mature adults at 35 dpe in adult fish. Bar = 200µm. M = Male, F = female with eggs. Arrows indicate the location of L2 larvae. **E)** A magnification of the box in panel D. Note the numerous immature eggs and L2 worms.

**Figure 5:**
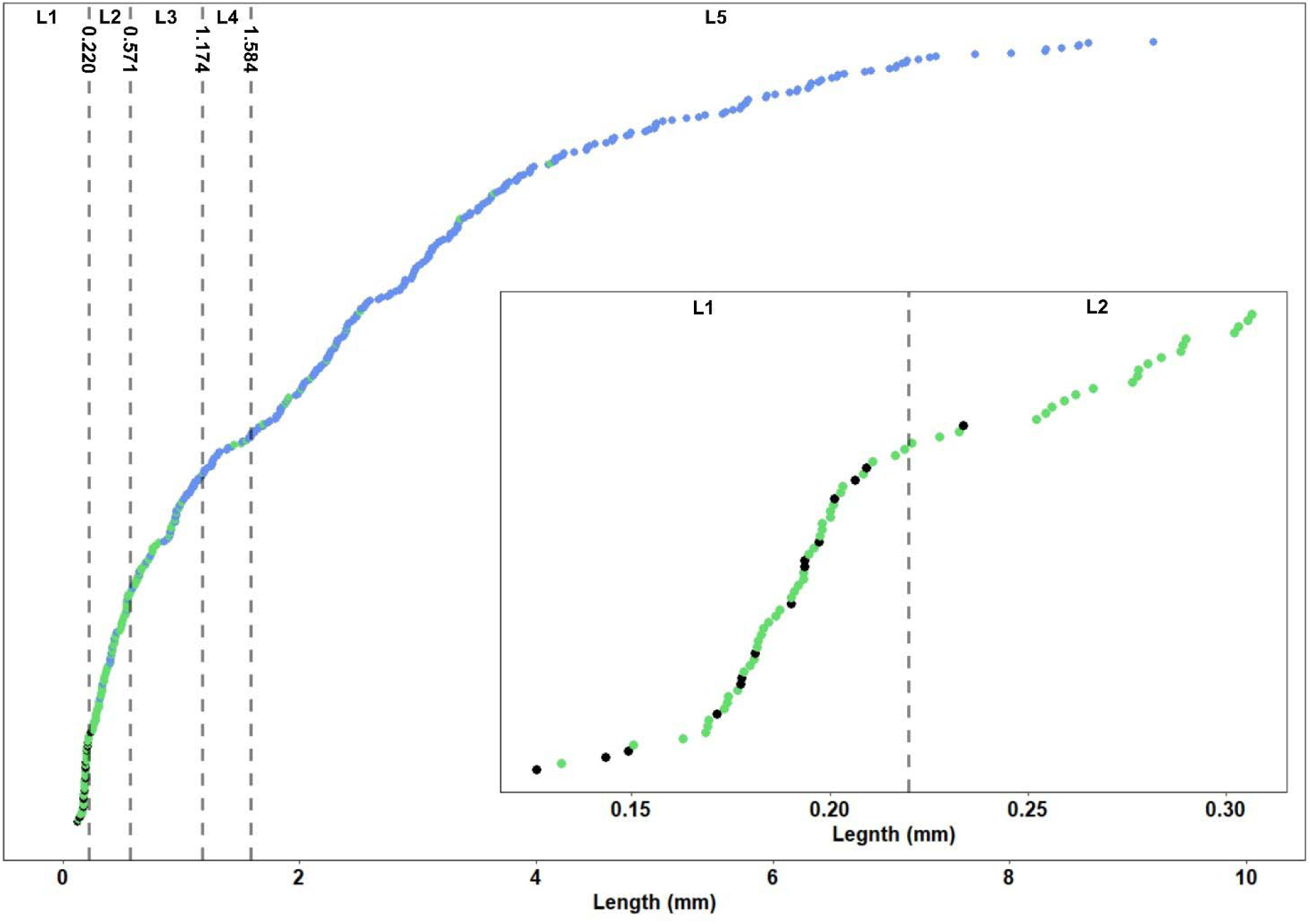
Worm size progression displayed in mm with change point analysis predictions of molting. Worms are ordered by length. Each dot represents an individual worm and dot color indicates the origin of data (Black dots = in vitro culture, green dots = larval fish, blue dots = adult fish). Lower right insert plot is enlarged region of early infection highlighting size location of worms from eggs (black) and those from larval fish (green). Note a few worms are unusually small and there is a clear gap between the predicted L1 and L2 change point.

**Figure 6:**
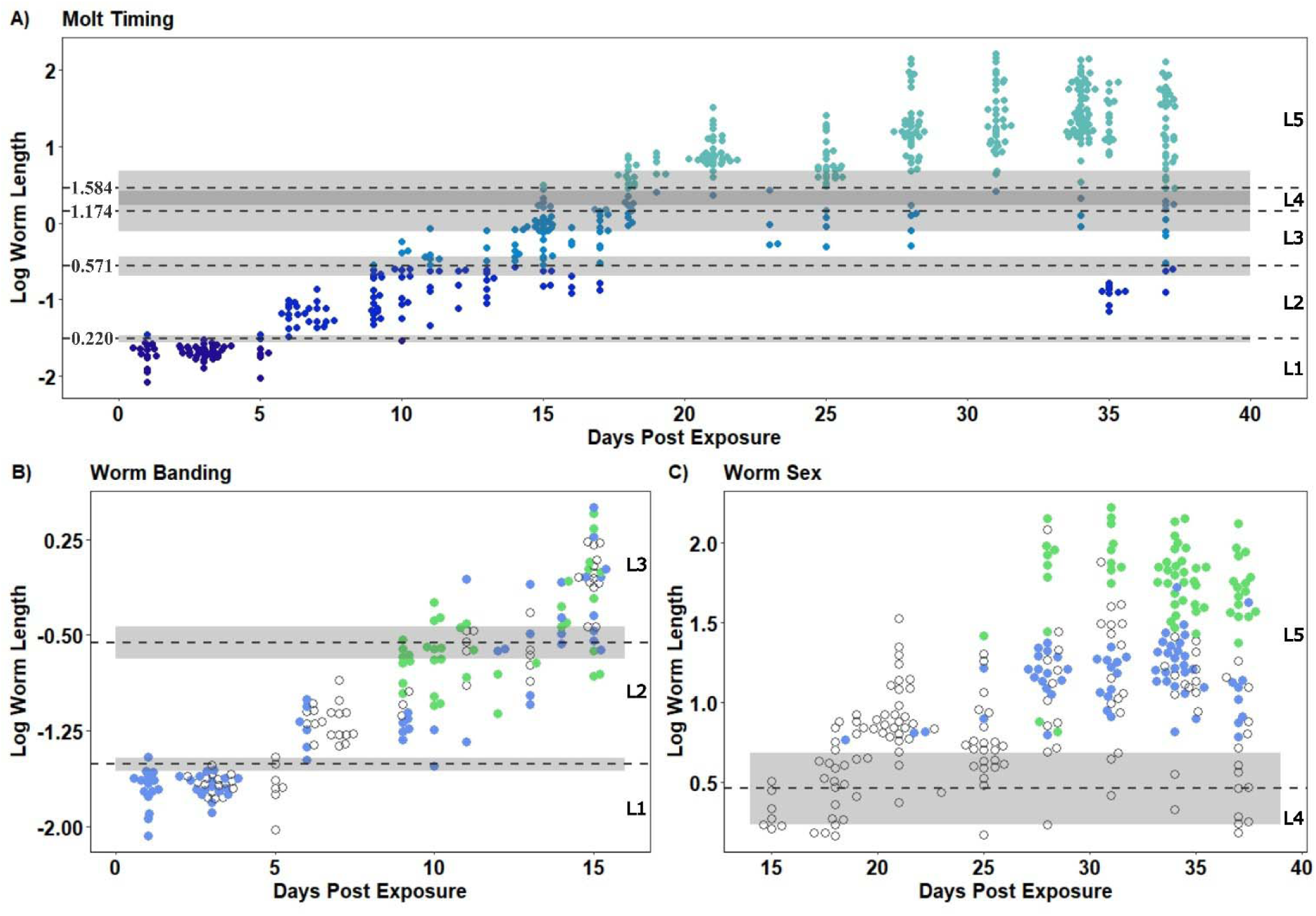
*P. tomentosa* development in larval and adult zebrafish between 1 and 37 days post exposure based on data from adult and larval fish exposures. **A)** The transition points (L1-L5) determined from the change point analysis. The Y axis is the log worm length with transition point size included. Each dot represents an individual worm, with each unique color representing developmental stages. The dashed lines represent the mean predicted change point for each molt threshold with the change point value in mm and the grey shaded areas represents 1 standard deviation. **B)** The appearance of worm banding with the same axis format as panel A. Open dots = unclassified worms, blue = unbanded worms, and green = worms with stichocyte bands. **C)** Later infection displays the sex of worms across the L4/L5 breakpoints using the same axis classification as panel A. Open=immature worms, blue = males, green = gravid females.

### 3.4. L1 Worm Phenotypes

The majority of the L1 worms were classified from *in vitro* cultures and larval fish between 3-5 dpe. This stage ranged in length from 0.126-0.219mm, with a mean size of 0.185mm. (Figs. 3a, 5,6). A total 14 of 15 worms from *in vitro* cultures and 39 worms from 3-5 dpe fish were classified as L1s. The shortest worm was from the *in vitro* group, but the next came from a 5 dpe larval fish, 0.126mm and 0.132mm respectively (Fig. 5). Overall, the five smallest worms appear to be distinctly shorter than the other L1s (Fig. 5) and would be considered as hypertrophic (runts). Many of the L1 worms originating from fish appeared to be straighter and more rigid in comparison to all worms from the *in vitro* cultures (Fig. 3A-C). Two worms were on the threshold of the L1/L2 change point, 1 *in vitro* worm classified as L2 and one larval worm at 10 dpe. These worms fell within the standard deviation of the L1/L2 change point.

### 3.5. L2-L4 Worm Phenotypes

L2 worms were first classified in 5 dpe fish and ranged from 0.220-0.564mm with a mean size of 0.392mm. Starting at 9 dpe some of the L2 worms expressed banded stichocytes, a trait not observed in L1 worms (Figs. 3 D-F, 6b.). No stylets were observed, as reported in L2 larvae as *Trichuris ovis* (Thapar and Singh 1964). L3s were designated starting in 9 dpe fish and ranged in length from 0.577-1.170mm with a mean size of 0.858 mm (Figs. 5,6). Aside from the increased length, there were no distinctive other morphological features. L4 worms were first detected in 15 dpe fish and ranged in length between 1.180-1.569mm with a mean size of 1.327, with no other additional morphological developments.

### 3.6. L5s and Worm Adult Phenotypes

The majority of L5 worms were mostly identified from adult fish and some from larval fish between 15-37 dpe. L5 worms from adult and larval fish ranged in size between 1.585-9.218mm with a mean size of 3.716 mm. L5 worms originating from larval fish (n=15) ranged in size between 1.693-4.125 mm (Fig. 5,6). At 25 dpe, two mature worms were observed in larval fish: a 4.125mm female with immature eggs (Fig. 6a) and a 3.359mm male (Trial Q). Males and females were sexually dimorphic with mature females being distinctly longer than males. Males ranged in size from 2.141-5.600mm, with the first male observed at 18 dpe (smallest male), and most of the males detected after 20 dpe. Females ranged in size between 2.255-9.218mm, with majority of females appearing at 34 dpe.

### 3.7. *P. tomentosa* Asynchronous Infections

Worm development exhibited some degree of asynchrony, particularly as the infection progressed. At 10 dpe, there was only one L1 (worms < 0.220mm) observed in all fish. In contrast, 15 to 18 dpe fish contained multiple developmental stages (L2-L5). Likewise, 25 dpe fish were infected with worms that ranged from L3 to sexually mature adults. At 35 and 37 dpe, L2 worms appeared again, suggesting a second cycle of infection.

### 3.8. Larval Fish Mortality

Using one trial (Q), we recorded mortality in exposed larval fish over 25 days (Fig. 7). The percent daily mortality was highest at 9-10 dpe (9.68%), and few fish survived past 15 dpe (Fig. 7). Larval fish mortality was linked to worm burden as the highest worm burdens were observed in fish before 10 dpe – i.e., fish assessed past 10 dpe had less than 4 worms/fish. At 9 and 10 dpe the mean worm lengths were compared between moribund and apparently healthy fish (Trial P). The mean worm size in healthy vs moribund fish were 0.373±0.085 vs 0.435±0.126mm at 9 dpe, and those in healthy vs moribund were 0.429±0.172 vs 0.563±0.145mm at 10 dpe. However, the size differences were statistically insignificant (p > 0.05), perhaps due to small sample sizes (9 dpe-14 fish, 10 dpe-13 fish).

**Figure 7:**
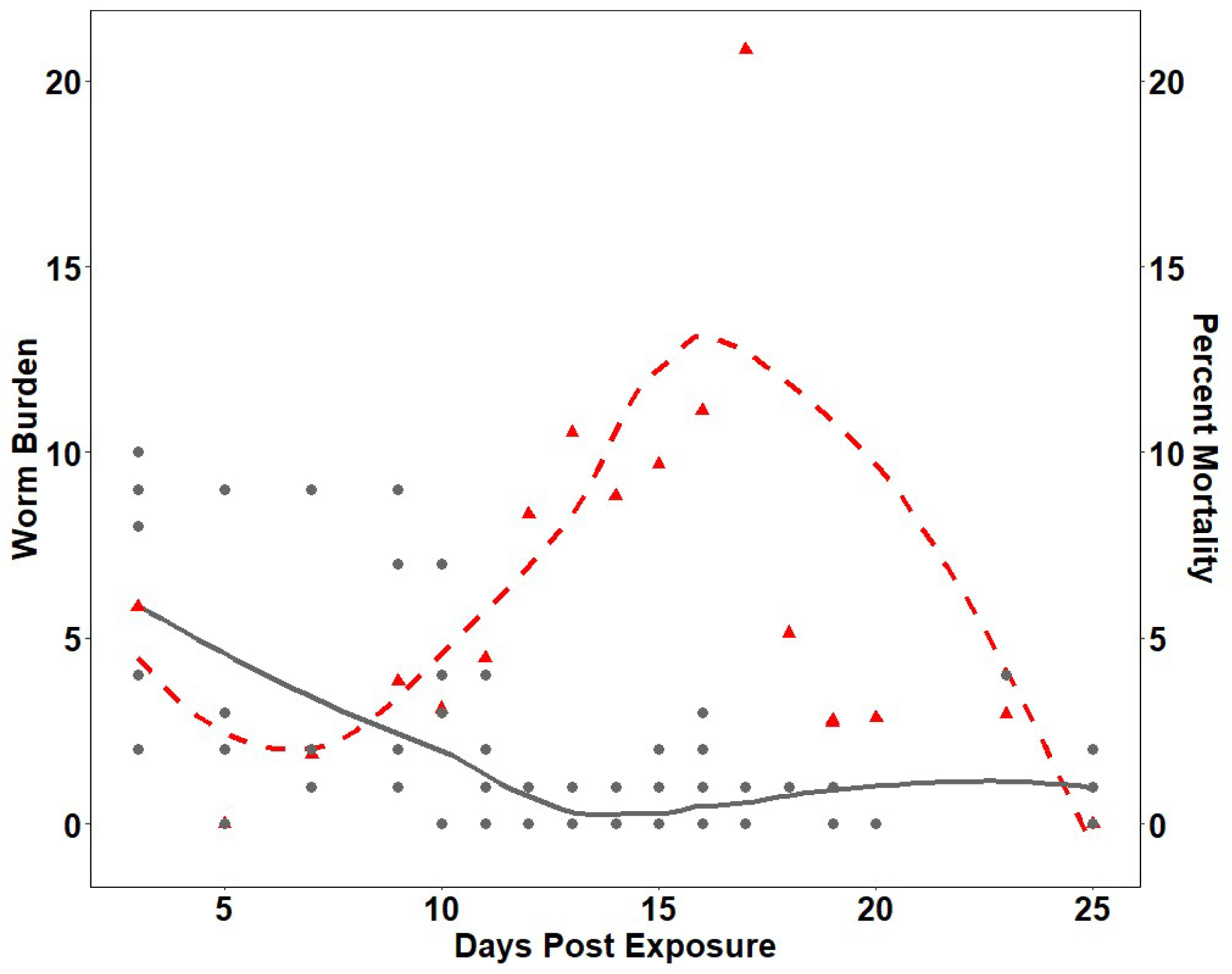
Parasite burden compared to mortality in larval fish in Trial Q. Red dashed line and triangles: daily percent mortality. Grey lines and circles: fish worm burden. Heavily infected fish die early, with mortality increasing after 8 days post exposure, most severe between 10-17 days, then decreasing after heavily infected fish died.

### 3.9. Emamectin Benzoate Exposure

The macrocyclic lactone emamectin benzoate was evaluated with the larval fish platform because this drug efficacious for treating infections in adult zebrafish (Collymore et al., 2014; Kent et al., 2019). Nontoxic concentrations were identified by aqueously exposing 10 and 28 dpf zebrafish in 24 well plates for 24hr. Fish at 28 dpf tolerated higher concentrations of emamectin benzoate than 10 dpf fish (SI3). 10 dpf fish had an LC50 0.08µM whereas 28dpf fish had an LC50 of 1.23µM. With this toxicity data, we exposed fish 0.025, 0.035, or 0.7µM of emamectin benzoate in our infection trials J, M, and N respectively. At 0.025µM of emamectin benzoate reduced infection, was not significant (p > 0.05, Wilcoxon rank sum test) (Fig 5a). However, 7/18 worms exposed to 0.025µM emamectin benzoate exhibited a distinct posterior vacuole (Fig 5b). In contrast only 1/18 worms in the unexposed group exhibited this vacuole (Fig. 5B). Trial M2 and N2 exposed 30 dpf fish to 0.035 or 0.7uM of emamectin benzoate, respectively and this drug reduced in the intensity of infection (p = 0.022, 0.032, respectively Wilcoxon rank sum test) (Fig. 8C, D). Also, in Trial N2, worm behavioral changes were noted in the emamectin benzoate exposed fish. Amongst the four worms seen in these treated fish (3/14 fish infected), one showed no motility, and another worm exhibited minor, stationary movement. In contrast, all 33 worms observed in untreated fish (8/14 fish infected), were very active, moving about in the intestine and actively probing with their anterior end.

**Figure 8:**
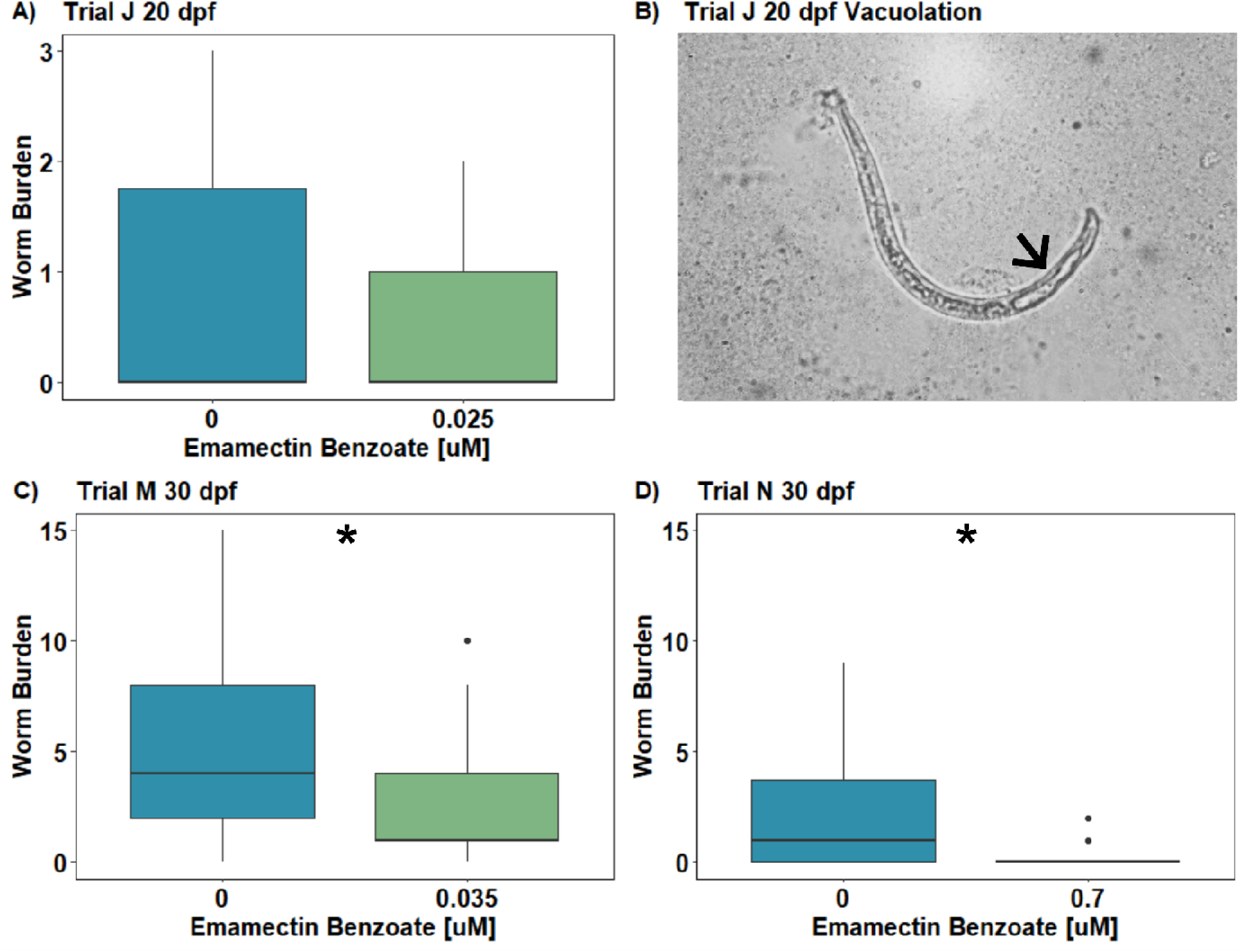
Emamectin benzoate efficacy with *P. tomentosa*. Zebrafish were exposed to 50 eggs/fish and compounds, and worm burdens were assessed at 3 days post exposure. Differences in abundance were evaluated using a Wilcox Ranks Sum Test (worm burden ∼ [emamectin benzoate]). **A)** Trial J, 20 dpf fish exposed to 0.025µM of emamectin benzoate had no significant difference in worm burden than unexposed fish, p > 0.05. **B)** An emamectin benzoate exposed worm from Trial J2, expresses vacuolization in the posterior region (arrow). **C)** Trial M. 30 dpf fish exposed to 0.035µM with significantly (*) lower abundance in treated fish p = 0.022. **D)** Trial N. 30 dpf fish exposed to 0.7µM of emamectin benzoate, with significantly (*) lower abundance in treated fish, p *=* 0.032.

## 4. DISCUSSION

The adult zebrafish/*P. tometosa* model has become a robust model used across several research avenues (Collymore et al., 2014; Gaulke et al., 2019; Hammer et al., 2024; Kent et al., 2019, 2002), and here we successfully translated it into a higher throughput larval fish platform for anthelmintic drug discovery. Our new assay includes infection exposures within 24 well plates and has been validated using the macrocyclic lactone emamectin benzoate, reducing the number of infected larval fish. This new platform benefits from increased sample sizes, shorter endpoint times for evaluating infections, a smaller amount of drug needed to test treatment effects, and fewer larvated eggs to establish reliable infections.

Our first experiments focused on selecting the optimal fish age for the platform. Late-stage larval fish (about 25-30 dpe) were ideal; they afforded all the advantages of larval fish in general, with the 24 well plate format, but showed more robust infections and less mortality than younger larval fish. Improvements to infections can be attributed to changes in the size of intestine, mouth size, and fish behavior. Fish at 5-8 dpf likely had low infection rates because of their smaller mouth. Previous studies have identified that the optimal particle size for early-stage zebrafish larvae feeding was 0.021– 0.045mm (Önal and Langdon, 2016), which is a bit smaller than *P. tomentosa* eggs at 0.065 by 0.030mm (length, width). In other words, the likelihood of 5-8 dpf zebrafish ingesting *P. tomentosa* eggs is reduced because the egg is simply too large for their mouth. Older larvae have a larger mouth; the optimum particle size for 15 dpf is 0.047–0.075mm size, closer to the size of the parasite eggs (Önal and Langdon, 2016). Moreover, these older fish actively swim and seek food, in contrast to early-stage fish that are stationary and only dart at food when it floats nearby (Parichy et al., 2009). As *P. tomentosa* eggs are stationary, and require water movement to be mobile, actively seeking food enhances the likelihood that a fish would consume an egg. Another benefit of older larvae is that both exposed and control fish survived better. Regarding the former, this is likely because their larger intestines and overall body size allow them to tolerate infections for longer durations.

Developing robust *in vivo* parasite models for drug discovery requires an in depth understanding of the biology and development of the parasite. Understanding *P. tomentosa* development is important for anthelmintic discovery because some drugs may have their most profound effects during sensitive developmental windows (e.g., transition through L1-L5 stages), induce morphologic changes, or alter worm sexual maturation and fecundity. However, little information exists about the development of *P. tomentosa* or the development of many genera of capillarid nematodes (Moravec et al., 1987). Many of the capillariids life cycles have not been completely studied (Moravec et al., 1987). Assessing worm development from *in vitro* egg hatching, larval and adult fish infections enabled us to document the entire development of *P. tomentosa* development from initial infection to maturation. Starting with eggs, *P. tomentosa* requires about 5-7 days at 28°C to larvate (Martins et al., 2017), which is similar to *Capillaria* (*Aonchotheca*) *philippiensis* in fish, whose eggs also develop in water (Cross and Basaca-Sevilla, 1991). *Baruscapillaria* spp. in birds also have a short larvation time, whereas most other terrestrial capillarids from mammals require several weeks to larvate (Moravec et al., 1987).

No molts or dramatic size breaks were observed after evaluating 488 worms across different developmental stages (Fig. 5). Therefore, a change point analysis (CPA) was used to estimate the timing and worm size of each molting event. This approach has been widely used in ecological studies to identify changes in populations and community composition (Thomson et al., 2010), but to our knowledge it has not been used to distinguish larval stages of nematodes. The biological relevance of our approach was supported by correlating changes in nematode morphology and the time order of our predictions. For example, banded stichocytes were only seen after L2s were observed and starting at 9 dpe (Fig. 8).

It is assumed that *P. tomentosa* undergoes ecdysis through 4 molt stages (i.e., L1-L5) as this is pattern in other capillarids and with nematodes in general (Anderson, 2000; Moravec et al., 1987). In fish host, *Capillaria philippinensis* develops to the L3 stage (Cross and Basaca-Sevilla, 1991). However, the most detailed descriptions of development through all various larval stages are the capillarids from certain birds (Moravec et al. 1987). Although birds have a higher body temperature than zebrafish as used here, *Baruscapillaria* spp. of domestic fowls have a similar timing of ecdysis to *P.tomentosa*. (Shlikas, 1965, 1966, 1967a, 1967b). The inclusion of larvae from hatched eggs provides a foundation for the size of early L1s. First hatched larvae range from about 0.13-0.16mm in length for *Baruscapillaria* spp., and then some L1 worms later grew to 0.200mm (Shlikas, 1967b). Early larval forms (L1s) of *C. philippinensis* in fish were also similar, 0.13-0.15mm (Cross and Basaca-Sevilla, 1991). Our designated L1 category contained majority of *in vitro* cultured worms and most worms from fish between 3-5 dpe. The lengths from these two sources were intermixed between 0.126-0.219mm, (Fig. 5). The L1s from our study showed two morphologies; more curved, slightly shorter worms were very similar to those from in vitro hatched eggs (Fig. 2A, B), and more ridged and straight worms (Fig 2c). This change in morphology could be attributed to worms inflating (taking on fluid) after hatching (O’Sullivan et al., 2020; Wright, 1975). There were a few unusually short worms (0.126-0.149mm), and the second shortest worm at 0.132mm was observed in a fish at 5 dpe. Previous studies have shown that high worm burdens create competition for resources resulting in stunted worms (Heins and Baker, 2011), but this very short worm occurred in a fish with only one worm.

The L1/L2 change point was predicted at 0.220mm, first seen at 3 dpe, except for one unusually long worm measured from an egg. L2s reached a maximum length of 0.564mm. Likewise, L2s of *B. obsignata* and *B. anseris* were similar in length and first occurred at 7 dpe (Shlikas, 1967b). In our study, L2s persisted for the most part until 17 dpe (Fig. 6). Shlikas (1967b) reported the first appearance of L2s at 8 days, but did not report how long L2s persisted. Stichocyte bands, a characteristic of *P. tomentosa* (Moravec, 2001), were first seen in L2s at 9 dpe and were observed throughout the remaining developmental stages. However, these bands were inconsistently observed, even in larger worms (e.g., L3 and L4 stages). In addition, an anterior stylet was not seen in either L1 or L2s as reported for *B. aneris* or *T. ovis*, respectively (Gobind and Suresh, 1954; Shlikas, 1967b). No L2s were observed between 18-34 dpe, indicating that all L2s had matured. However, a few L2 size worms were seen at 35 and 37 dpe, which was likely due to a second cycle of infection as L2s. This is in accordance with previous studies where eggs were not released from females until about 20-25 dpe (Kent et al., 2018).

Worms categorized as L3 (0.577-1.170 mm) were first predicted to occur at 9 dpe and L4s at 15 dpe (1.18-1.569mm). Likewise, in *Baruscapillaria* species L3 and L4 developmental stages are observed at 10 and 13 dpe, respectively (Shlikas, 1967b). Regarding lengths, L3s of *Baruscapillarias* spp., they are 0.8-2.2mm for *B. obsignata* and up to 2.75mm for *B. anseris,* (Shlikas, 1967b), somewhat longer than our predicated length range for L4s in our study *P. tomentosa.* We found that both L3 and L4 developmental stages were asynchronous in their appearance as multiple developmental stages were observed at the same time and often in the same fish.

Our predicted L3/L4 change point was 1.174mm. Moravec (1983) conducted a field survey of *P. tomentosa* in carp (*Cyprinus carpio*) and tench (*Tinca tinca*), and the shortest worms observed were 1.42mm, which would be in our L4 range. Moravec (1983) identified these worms as immature females by the presence of a small vulva primordium, but we did not detect this structure in our study. Whereas our studies with zebrafish and those by others with other fishes (Lomakin and Trofimenko 1982) verified that that *P. tomentosa* can be directly with larvated eggs, oligochaetes may serve as paratenic hosts (Lomakki and Trofinmenko 1982; Moravec 2011). Stages consistent with our designations of L1-L3 not observed in the carp intestines from the field, and hence Moravec (1983) suggested that paratenic host played a significant role in the life cycle of the parasite in the environment. However, we discovered that even in the relatively small intestines of zebrafish compared to that of carp, it is difficult to detect early infections in wet mounts (Schuster et al. 2023).

L5 worms were first observed at 15 dpe, with a predicted edysis size of 1.585mm. The considerable size differences amongst worms in the L5/adult category was attributed to the sexual dimorphism between males and females, which is observed is the case with other reports for *P. tomentosa* and other *Pseudocapillaria* species (Kent et al., 2002; Moravec, 2001). Males were identified by the presence of a spicule, and the first males seen at 18 dpe and they ranged in length from 2.141-5.600mm, all longer than the predicted change point for L4/L5.

Mature females were distinguished by the presence of eggs and measured 2.255-9.218mm in length. Female genitalia (e.g., vulva primordia) were not noted in worms without eggs, although a few large worms without eggs occurred within the designated female range. The identification of females in our study was based on the presence of eggs, which must occur after males are present and worms have mated. Indeed, we observed males as early as 18 dpe, while all but one gravid females were first not seen until 28 dpe. The first appearance of gravid females in this study agrees with previous studies which first observed gravid female worms, first observed at about between 21-28 dpe (Collymore et al., 2014; Gaulke et al., 2019, 2016; Hammer et al., 2024; Kent et al., 2002; Sieler et al., 2025).

The prepatency time with *P. tomentosa* is shorter than many capillarids, except in vertebrate hosts that first become by feeding on an intermediate host –i.e., are first infected with L2 or L3s, rather than larvated eggs (Moravec et al., 1987). For example, with *Capillaria pterophylli* of fish with a direct life cycle eggs are not seen until about 90 days post infection at 20-23°C (Moravec et al., 1987). A few terrestrial capillilards of birds in the genera *Aonchotheca* and *Baruscapillaria* have a comparable prepatency times to *P. tomentosa*, whereas capillarids of mammals (e.g., *Pearsonema mucronatea*) and *Trichuris* spp require more time (Gobind and Suresh, 1954; Moravec et al., 1987; Shlikas, 1967b). For example, it was reported that whipworms often require 2-4 months before females become gravid and release eggs (Hurst and Else, 2013). It is interesting that *P. tomentosa* at 28°C develops faster than many capillarid species in mammals, with their much higher body temperatures.

Over dispersion in macroparasites is essentially a law in parasitology (Poulin, 2007), and this extends to *P. tomentosa* in zebrafish (Kent et al., 2018). Overall, the infections between adult and larval fish were similar. Although social interactions and host hierarchies may affect parasite burdens, negative binomial distributions of parasites were also seen in adult zebrafish that were infected in isolation (Hammer et al., 2024) as well as in larval fish in our study. Another important aspect of comparing our fish models to mammals (i.e., whipworms) is the location and pathogenesis of infection. In the adult zebrafish infection model worms penetrate the epithelium and cause profound lesions (Gaulke et al., 2019). Infection location and associated pathologic changes were similar in our larval fish at 3 dpe, with worms embedded in the epithelium and associated proliferative changes at this location.

While growing out infected fish, adult fish survived well throughout the infection, whereas larval fish survival was poor between 10-17 dpe. As seen with early versus late-stage larval fish, poor survival can be attributed to the smaller gut size and the overall size of the larvae compared to adults. The intensity and size of worm infection influenced larval fish survival outcomes. Fish with high burdens (>5) did not persist past 10 dpe, then fish with low worm burdens mostly succumbed about 4-5 days later. Only a few infected larval fish survived past 20 dpe because worms increase in size, regardless of the intensity of infection, so only two mature worms were observed in larval fish. The worm sizes from the larval and adult models overlapped suggesting the concurrency between models. Hence, combining the adult fish assay with the larval version enabled assessment of entirety of *P. tomentosa* development.

The efficacy of emamectin benzoate in 28 dpf fish reflected our results with adult fish (Collymore et al., 2014; Kent et al., 2019). Adult and 28 dpf fish can tolerate similar emamectin benzoate concentrations demonstrating that older larval fish can be suitable alternative models for drug discovery (Kent et al., 2019). Furthermore, emamectin benzoate was more toxic to 10 dpf fish than 28 dpf fish, which is observed in mammalian species due to differences in biological processes (Scheuplein et al., 2002). The efficacy of emamectin benzoate in the larval fish platform was concentration dependent. At the lowest concentrations, vacuolated worms were detected with no reduction in infection intensity. In contrast, higher concentrations reduced infection intensity with no vacuolated worms. Changes in nematode morphology at low concentrations highlights new potential endpoints for this platform. Nematode vacuolation may be an appropriate endpoint because vacuolization has been reported in the pine wood nematode *Bursaphelenchus xylophilus* exposed to 7-indoindole or spectinabilin (Liu et al., 2019; Rajasekharan and Lee, 2020).

Integrating the larval zebrafish platform, along with the adult fish version when appropriate, into anthelmintic drug development pipelines provides a novel opportunity to enhance anthelmintic discovery. In comparison to our more established adult zebrafish platform, the larval zebrafish platform has some distinct advantages in assay throughput. Transitioning from 9L tanks to 24 well plates reduces the amount of labor and required resources. Smaller water volumes require less amounts of chemicals, which is particularly useful for limited or costly chemicals. Multi-well plates allow fish to be individually exposed to defined numbers of eggs for the entirety of the experiments, which enhances the assay’s sensitivity and statistical power. The larval fish platform also increases the speed of the assay readouts because endpoints can be assessed at 3 days in comparison to 3-4wk with adult fish without dissecting the intestine. Previous studies with adult fish, have found inconsistent detection and enumeration of worms by wet mounts before 15 dpe (Kent et al., 2018; Schuster et al., 2023). In contrast, L1s and L2 size worms, which occur shortly after exposure, can be more easily detected in wet mounts of larval fish because they have a more translucent and thinner walled intestines. Finaly, infections in 30 dpf fish can be evaluated faster because careful dissection of the intestine is not required.

In early-stage drug discovery, drug leads are prioritized using *in vitro* assays like *Caenorhabditis elegans* or filarial worms because these assays are rapid, flexible, and high throughput (Storey et al., 2014). However, many drug leads fail in subsequent clinical tests because they do not elucidate complex host drug interactions and toxicity. For example, the efficacy of ivermectin against heartworm *Dirofiliaria immitis* would not have been identified in *in vitro* approaches because ivermectin at the low concentrations for treating this worm required participation of the host immune system (Carithers, 2017; Vatta et al., 2014). Hence, *in vivo* assays expand opportunities to detect efficacious drugs.

Another attribute of both the larval and adult fish platforms is that the entire worm burden in the intestine is enumerated as the entire intestine is examined in wet mounts. In contrast, other *in vivo* evaluations usually extrapolate infection intensity from parasites in fecal samples. Zebrafish platforms can also assess other non-lethal endpoints, like worm growth and sexual development, as well as egg production. Zebrafish assay readouts are likely translatable to mammalian parasite species because it has already been shown that macrocyclic lactones (emamectin benzoate) and benzimidazoles (fendbendazole) are effective across major nematode taxa, including *P. tomentosa* (Collymore et al., 2014; Kent et al., 2019; Maley et al., 2013).

Nematode drug resistance is rising, and new tools are required to improve drug lead prioritization. The zebrafish infection model bridges the gap between *in vitro* and *in vivo* testing, enhancing drug lead detection. This zebrafish model is an ideal intermediate model before moving into mammalian models because they are high throughput and account for host-parasite interactions, drug metabolism, and toxicity, while maintaining a high level of sensitivity to a diverse suite of endpoints. We are now moving forward with both the larval and adult fish platforms to screen a library of compounds that negatively correlated with infection intensity (Hammer et al. 2024).

## Data Statement

The metadata generated for this manuscript is available at the GetHub repository using the following url: https://github.com/ConnorLeong/Pseudocapillaria_tomentosa_Development.

## Credit authorship contribution statement

**Connor Leong:** Conceptualization, Methodology, Investigation, Formal Analysis, Visualization, Writing Original Draft, Writing – review and editing. **Ruby Scanlon:** Investigation, Writing review and editing. **Aisling Kyne:** Investigation, Writing review and editing **Thomas J Sharpton:** Conceptualization, Supervision, Formal Analysis, Funding Acquisition, Writing Original Draft, Writing – review and editing. **Michael L Kent:** Conceptualization, Supervision, Formal Analysis, Funding Acquisition, Writing Original Draft, Writing – review and editing.

## Acknowledgements

We would like to thank Ryan Lopez and the staff at the Sinnhuber Aquatic Research Laboratory, Oregon State University for providing the zebrafish. We also thank Dr. James Peterson, Oregon Cooperative Fish and Wildlife Research Unit, U.S. Geological Survey for recommendations with change point analysis. This research was supported by the National Science Foundation S2244E and Agricultural Research Foundation, Oregon State University HT038B and HT038C. Provision of fish for this publication was provided by the Sinnhuber Aquatic Research Laboratory, which is supported by the National Institute of Environmental Health Sciences of the National Institutes of Health under Award Number 30ES030287. The content is solely the responsibility of the authors and does not necessarily represent the official views of the National Institutes of Health. None of the authors have any competing interest.

