## Supplemental figures for "*Pseudocapillaria tomentosa* Infections in Laboratory Larval and Adult Zebrafish (*Danio rerio*): Development and Advances in an *In Vivo* Anthelmintic Drug Discovery Model"

**Appendix A. Supplementary Information**

The following information is the supplementary material for the manuscript: *Pseudocapillaria tomentosa* Infections in Laboratory Larval and Adult Zebrafish (*Danio rerio*): Development and Advances in an *In Vivo* Anthelmintic Drug Discovery Model

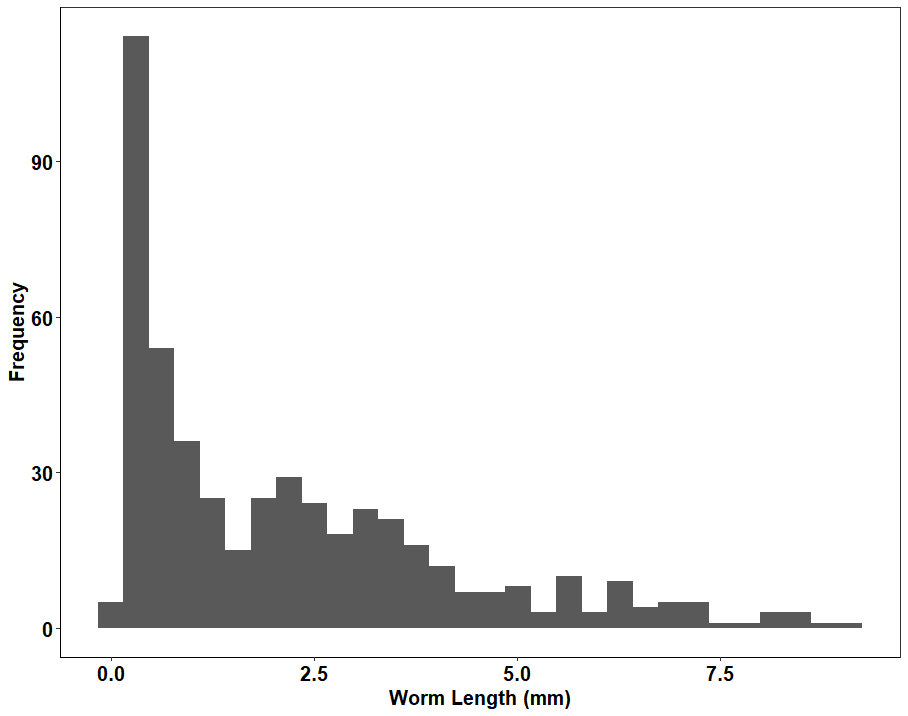

**SI.1:** The distribution of worm lengths from 1 to 37 dpe. Worm lengths were obtained from 30 dpf infected zebrafish and hatched eggs from *in vitro* cultures. Worm length in mm is indicated on the x-axis and the frequency of worms is indicated on the y-axis.

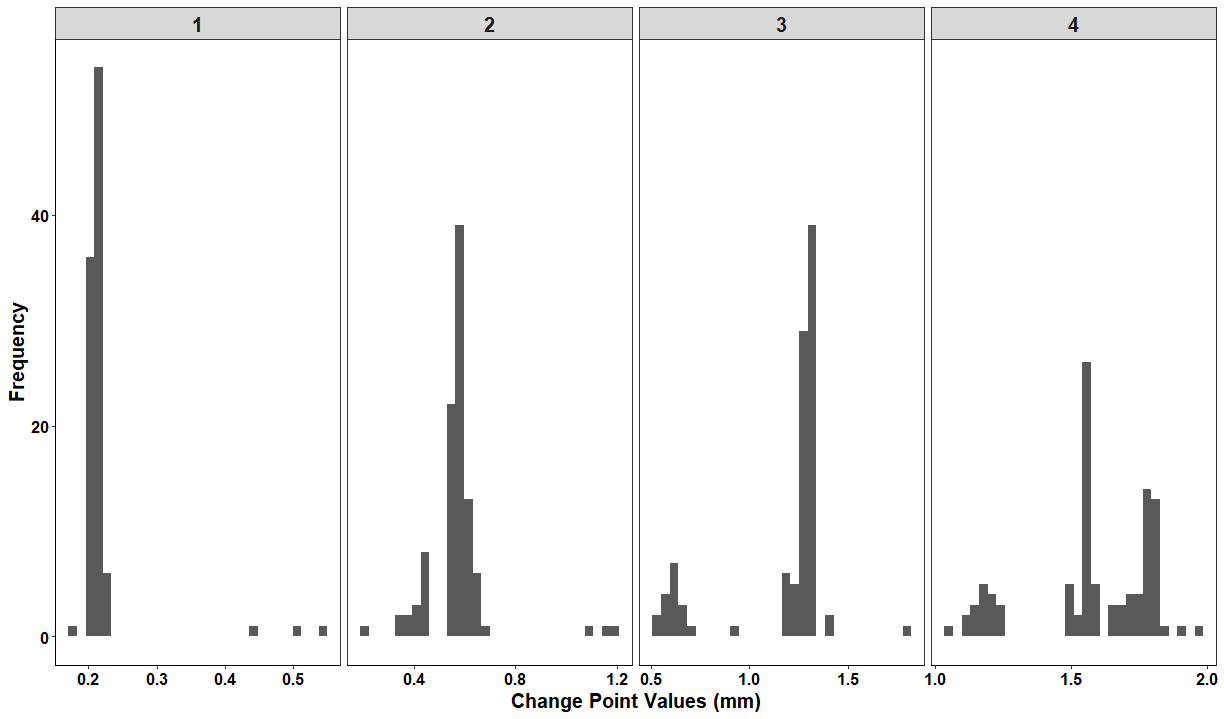

**SI.2:** The frequency of individual change point values for all change point analysis bootstrap iterations (100xs). The four panels represent change point 1-4 respectively. Within each panel the x-axis represents the change point values in mm and y-axis represents their frequency.

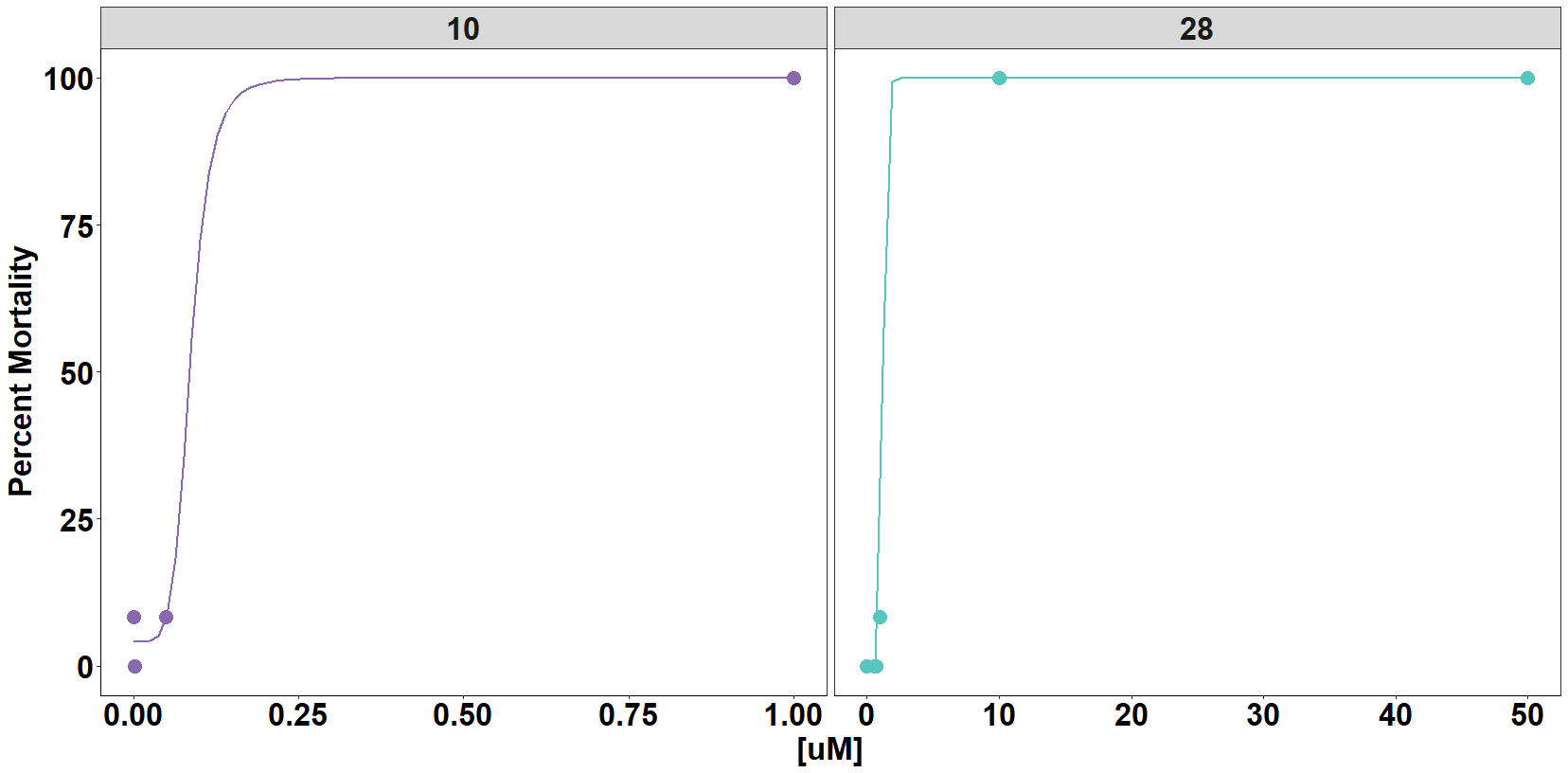

**SI.3:** The emamectin benzoate toxicity assessments for 10 and 28 dpf. The panel labels and line colors represent the fish age (purple = 10, turquoise = 28). Each point represents the specific emamectin benzoate concentration tested on the X axis and the corresponding percent mortality on the (y-axis).
